# Computational simulations reproduce *in vivo* population receptive field mapping fMRI results

**DOI:** 10.64898/2026.07.14.738461

**Authors:** Siddharth Mittal, Michael Woletz, David Linhardt, Christian Windischberger

**Affiliations:** High field MR Center, Center for Medical Physics and Biomedical Engineering, Medical University of Vienna, Vienna, Austria

**Keywords:** fMRI, retinotopy, population receptive field, simulation, visual neuroscience

## Abstract

Population receptive field (pRF) mapping is widely used to characterize retinotopic organization based on functional magnetic resonance imaging (fMRI) data. Despite its broad adoption, the factors governing intra- and inter-subject variability in pRF estimates remain incompletely understood, limiting the ability to evaluate and optimize visual stimulation paradigms prior to data collection. Here, we investigate whether large-scale simulations can reproduce *in vivo* run-to-run variability patterns observed in pRF mapping and provide mechanistic insight into their origins. With GEMSim-pRF, our newly proposed computational framework for large-scale simulation and estimation of pRF responses, we generated millions of synthetic fMRI time courses across a wide range of receptive field parameters and noise conditions. We analyzed the variability of pRF estimation results derived from simulations and compared them with *in vivo* data from the publicly available NYU Retinotopy Dataset. Here we show that our simulation results matched the characteristic eccentricity-dependent variability observed in empirical pRF estimates. These findings show that key variability patterns observed in empirical pRF mapping can be successfully reproduced in large-scale simulations, establishing simulation-based analysis as a practical approach for understanding, predicting, evaluating and ultimately improving the behaviour of retinotopic mapping paradigms before empirical data collection.

## 1 Introduction

Introduced in the late 2000s, population receptive field (pRF)-based mapping [Dumoulin and Wandell, 2008] has become the prime method for determining retinotopic associations between the visual field and the visual cortex. This approach is based on the acquisition of *in vivo* functional magnetic resonance imaging (fMRI) data to determine neural activation patterns in the brain. During the fMRI scan, participants are presented with specific visual stimulus patterns that typically shift throughout the visual field. Retinotopic mapping results are obtained by analysing the fMRI data using pRF mapping software packages such as mrVista [Dumoulin and Wandell, 2008], analyzePRF [Kay et al., 2013], popeye [DeSimone et al., 2016], SamSrfX [Schwarzkopf, 2018], Braincoder [de Hollander et al., 2024], or the very recently introduced GPU-based software GEM-pRF [Mittal et al., 2026]. Typically, these pRF mapping tools relate an empirically measured fMRI time course to a pRF defined by its specific location and size in the visual field. Although the specific parameter estimation strategies differ across software implementations, the underlying principle is shared: pRF parameters are estimated by determining the pRF parameter configuration with the best match between predicted and observed fMRI signal. As described in the existing literature, the predicted time courses depend on the pRF model used. Typical approaches include two-dimensional Gaussian modeling [Dumoulin and Wandell, 2008], difference-of-Gaussians modeling [Zuiderbaan et al., 2012], or compressive spatial summation modeling [Kay et al., 2013].

For adequate interpretation of retinotopic mapping results, particularly in clinical and translational applications, it is essential to understand not only the pRF estimates themselves but also any bias introduced by the spatiotemporal properties of visual stimulation paradigms themselves. For example, [Prabhakaran et al., 2021] showed that fMRI-based visual field reconstructions depend strongly on the visual stimulation paradigm, with different approaches exhibiting distinct sensitivity-specificity trade-offs. Similarly, several studies have reported systematic stimulus-dependent differences in pRF size, eccentricity, visual field coverage, and model fit quality [Alvarez et al., 2015, Chang et al., 2025a, Linhardt et al., 2021]. In order to compare different analysis pipelines, Lerma-Usabiaga et al. [2020] introduced a powerful validation framework for assessing parameter recovery under controlled noise and modeling assumptions. Together, these findings indicate that the variability in pRF estimates is not solely determined by measurement noise, but also emerges from interactions between stimulus design, model assumptions, and the estimation procedure itself.

Understanding the origins of this variability remains challenging using empirical data alone. While repeated measurements can provide estimates of run-to-run variability, the number of repetitions that can be acquired within a single participant is inherently limited by scan duration, participant burden, and experimental cost. Consequently, empirical studies are often statistically underpowered for systematically investigating how variability changes across visual field location, pRF size, noise level, or stimulus design. Existing large-scale retinotopy datasets further illustrate this limitation. The Human Connectome Project [Benson et al., 2018] provides multiple retinotopic stimuli across a large cohort but typically only a single run per stimulus. Conversely, the NYU retinotopy dataset [Himmelberg et al., 2021a] provides extensive repeated measurements but is restricted to a single stimulation paradigm. Similarly, the CHN retinotopy dataset [Chang et al., 2025b] includes multiple paradigms and sessions, but only a limited number of effective repetitions per stimulus configuration. As a result, empirical datasets alone provide limited opportunities to systematically characterize the mechanisms governing pRF estimation variability.

Simulations provide a potential solution to this problem as they could provide a powerful framework for understanding the origins of pRF estimation variability and for evaluating retinotopic stimulation paradigms prior to empirical data collection. By generating synthetic fMRI time courses with known ground-truth parameters, simulations make it possible to isolate and manipulate individual factors that influence pRF estimation behaviour. Computational constraints, however, limit the practical scale at which repeated pRF estimation can be performed.

To overcome these challenges, we are introducing GEMSim-pRF (GPU-Empowered Mapping & Simulations of pRFs), a framework that combines large-scale fMRI simulation with GPU-accelerated pRF estimation. By taking advantage of the GPU-accelerated performance of GEM-pRF [Mittal et al., 2026], the GEMSim-pRF framework enables millions of simulated pRF estimations across visual field locations, pRF sizes, and noise conditions. Using this framework, we herein investigate whether empirical variability can be reproduced in simulations and examine the factors that contribute to their emergence. Specifically, we compare the variability patterns derived from large-scale simulations with repeated-measurement variability observed in the NYU retinotopy dataset [Himmelberg et al., 2021a]. We show that the characteristic increase in variability with eccentricity emerges naturally in simulations and that a simple white Gaussian noise model is sufficient to reproduce the major empirical patterns. These findings establish large-scale simulation as a practical approach for understanding and predicting variability in pRF mapping and provide a foundation for future simulation-driven evaluation of retinotopic stimulation paradigms.

## 2 Methods

### 2.1 Empirical retinotopy data

To evaluate the correspondence between simulated and empirical measurements, we used the publicly available New York University (NYU) retinotopy dataset [Himmelberg et al., 2021a]. The dataset comprises functional MRI data from 44 healthy participants who completed a standard bar-aperture retinotopy experiment.

Data were acquired at the NYU Center for Brain Imaging using a 3T Siemens MAGNETOM Prisma scanner with a 64-channel head coil. Structural images consisted of T1-weighted scans, with additional T2-weighted images available for a subset (n=11) of participants. Functional data were collected using a multiband T2*-weighted EPI sequence (TR = 1 s, 2 mm isotropic resolution), along with reverse phase-encoded spin-echo images to enable distortion correction. All preprocessing was performed by the dataset authors using fMRIPrep [Esteban et al., 2019]. Further details on acquisition and preprocessing can be found in Himmelberg et al. [2021b].

### 2.2 Visual stimulus

The visual stimulation protocol used in this study corresponds to the bar-aperture paradigm employed in the New York University (NYU) retinotopy dataset [Himmelberg et al., 2021b]. This stimulus was used both for the empirical data and for the generation of simulated time courses, ensuring consistency between empirical and simulated analyses.

For the *in vivo* data collection, participants viewed dynamic visual content presented within a moving bar aperture (width 3.1°). The aperture contained a stimulus pattern of colorful objects, faces, and scenes of varying scales, which appeared randomly on a pink-noise background. Stimuli were confined to a circular aperture of 24.8° diameter. The bar traversed the visual field in eight sweeps, each consisting of 24 discrete steps presented at 1 s intervals, synchronized with MR image acquisition (TR = 1 s). Each scan lasted 192 s, and between 4 and 12 runs were acquired per participant.

For the generation of simulated fMRI time courses and for pRF estimation in both empirical and simulated data, the stimulus representation was reduced to a binary aperture sequence, which served as input for generating pRF model time courses. This abstraction isolates the stimulus geometry and timing, while omitting higher-order visual features that are not covered in pRF estimation.

### 2.3 pRF estimations

Population receptive field (pRF) parameters were estimated using GEM-pRF [Mittal et al., 2026], a GPU-accelerated software package (gemprf-0.1.11, https://pypi.org/project/gemprf/) for large-scale pRF estimations. The same estimation pipeline was applied consistently to both simulated and empirical datasets. GEM-pRF was selected because it combines high computational efficiency with estimation accuracy comparable to established pRF analysis tools, such as mrVista [Dumoulin and Wandell, 2008], making it well-suited for the large number of simulations required in the present framework. A dense sampling space was specified to ensure high-resolution parameter estimation. The search space employed a 219 × 219 spatial grid up to 37° diameter, resulting in a spacing of approximately 0.17° between adjacent grid points. pRF size (*σ*) was sampled using 24 linearly spaced values ranging from 0.1° to 3.0°. In this study, only grid-based pRF estimation was performed. As described in the original GEM-pRF framework, sufficiently dense grid sampling provides accurate estimates, while refinement primarily interpolates between neighboring grid points [Mittal et al., 2026]. Given the focus of this work is on evaluating overall variability trends and RMS error distributions rather than fine-grained parameter optimization, dense grid-based fitting was sufficient and substantially reduced computational overhead.

The pRFs were modeled as two-dimensional isotropic Gaussians. HRF parameters were specified to select a canonical double gamma SPM-style hemodynamic response function [Friston et al., 1998], which was normalized to have unit sum. Low-frequency temporal drifts were modeled using discrete cosine transform (DCT) basis functions, including a constant term and additional low-frequency components (0.5 and 1.0 cycles per scan). In order to keep the residual degrees-of-freedom between the simulated and empirical data consistent, no further subject-specific regressors were used for the empirical data analysis.

### 2.4 Simulated fMRI time courses

An overview of the complete GEMSim-pRF simulation framework is illustrated in Figure 1. GEMSim-pRF generates simulated fMRI time courses by adding noise to model predictions *p*(*t*; ***<u>θ</u>***), where ***<u>θ</u>*** denotes the ground-truth parameters of a simulated pRF. In the current framework, a common set of *N* noise realizations is generated and subsequently added to every simulated noise-free model time course. Based on the configuration used in the present study, *N* = 2000 noisy time courses were generated using white Gaussian noise sampled from 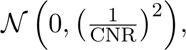 where CNR denotes the contrast-to-noise ratio.

**Figure 1:**
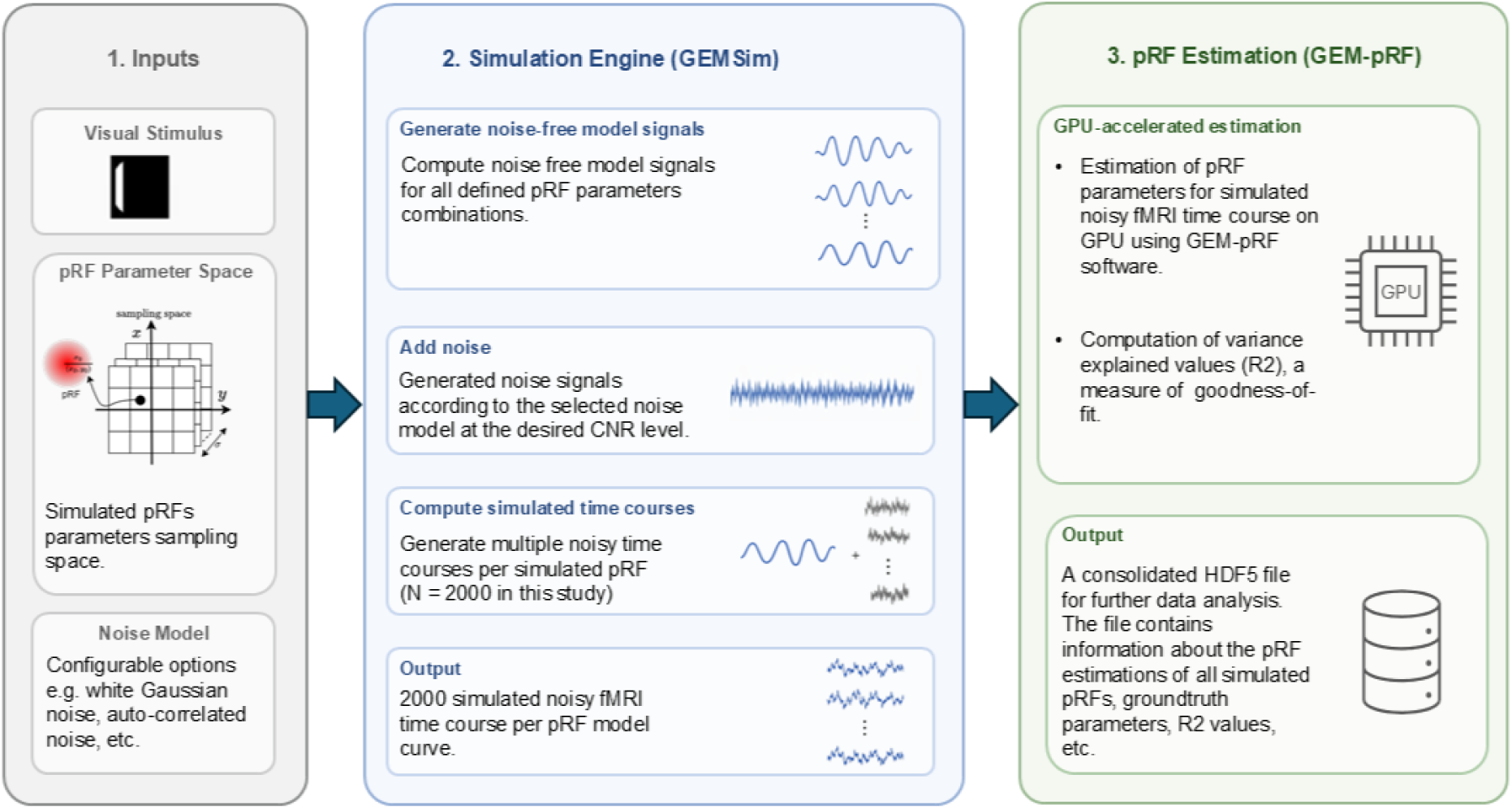
Overview of the GEMSim-pRF framework.

#### 2.4.1 Normalization factor

Our definition of the CNR assumes a unit-height contrast. We therefore normalize the response of each pRF to produce a unit-height response, i.e. the response will be equal to one for a fully stimulated pRF. Taking the regular sampling of the pRF into account and using the Riemann sum as numerical approximation, the normalization factor is therefore the infinite 2D integral of each pRF divided by the sampling spacing in *x* and *y*, Δ*x* and Δ*y* respectively:

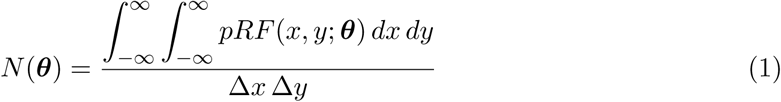

This model assumes a linear relationship for the input stimulus and the neural response and assumes that each simulated pRF will lead to the same response height for a complete stimulation.

#### 2.4.2 Computing CNR range from empirical data

Simulations were performed at representative CNR conditions selected from the range of CNR values observed in the empirical data. Only the fMRI time courses from the primary visual cortex (V1) of empirical data were used in this analysis. CNR values were computed individually for each empirical fMRI time course *y*(*t*). No *R*^2^ thresholds were applied when computing the empirical CNR distribution because such thresholds preferentially remove low-CNR vertices and would therefore bias the observed distribution toward higher CNR values. The resulting distribution was subsequently used to select representative simulation CNR levels.

For an estimated empirical pRF parameters ***θ̂***, the corresponding predicted time course *p*(*t*; ***θ̂***) was normalized using the normalization factor formulation described in Section 2.4.1 to ensure comparable signal amplitudes across vertices:

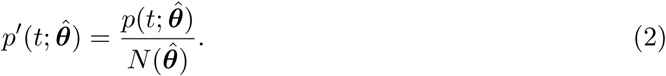

The normalized model time course *p^′^*(*t*; ***θ̂***) was used within a general linear model (GLM) framework to compute the residual error *ɛ*(*t*):

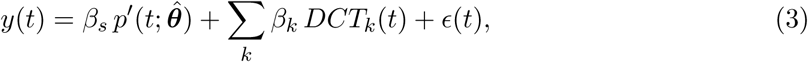

where *β_s_* represents the amplitude of the stimulus-driven response, *β_k_* are nuisance regressors capturing low-frequency temporal drifts via DCT basis functions.

As the scaling coefficient *β_s_* reflects a direct estimate of signal amplitude independent of nuisance regressors, the residuals need to be normalized to express noise relative to signal strength and ensure comparability across vertices using,

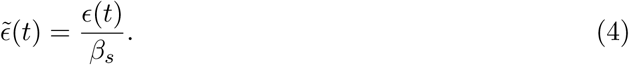

This results in the contrast-to-noise ratio as:

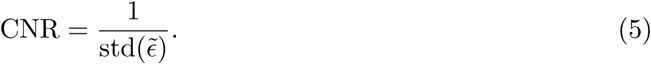

#### 2.4.3 Adding noise to generate simulated fMRI time courses

For a given CNR value, synthetic noise was generated and added to the normalized noise-free model time courses to produce the final simulated fMRI signals. For each simulated parameter set ***<u>θ</u>***, the corresponding normalized model prediction *p^′^*(*t*; ***<u>θ</u>***) was combined with noise realizations *ɛ*^(^*^n^*^)^(*t*) to generate simulated time courses:

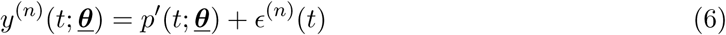

where *n* = 1*, . . ., N* denotes the individual noise realisations. The noise term *ɛ*^(^*^n^*^)^(*t*) was scaled to match the desired CNR levels derived from empirical data, ensuring that simulated time courses span realistic noise regimes while maintaining controlled and consistent signal scaling across all pRF parameter configurations.

### 2.5 Characterisation of pRF estimation variability

A series of analyses were designed to evaluate the extent to which simulations reproduce the variability observed in empirical measurements and to investigate the contribution of individual factors to pRF estimation variability under controlled simulation conditions.

#### 2.5.1 Variability at representative locations

To illustrate stimulus-dependent variability in a controlled setting, two representative pRF locations were selected: one at the center of the visual field and one at a peripheral location. For both cases, the pRF size was fixed to *σ*=1° to isolate the effect of spatial position on estimation variability.

For each location, 2000 noisy time courses were generated at the chosen CNR levels. The pRF parameters were then estimated independently for each time course. The resulting estimates (x,y) were visualized as scatter distributions to characterize the spatial dispersion of estimated pRF positions relative to the ground truth.

#### 2.5.2 Variability correspondence between empirical and simulated data

To assess whether variability observed in simulations reflects empirical trends, we compared the spatial spread of pRF estimates as a function of eccentricity between simulated and empirical datasets. The empirical analysis was performed using the pRF estimates from the primary visual cortex (V1) region of the NYU Retinotopy Dataset. Only pRF estimates with a variance explained (*R*^2^) of at least 10% were included in both empirical and simulated analyses. Variability was quantified using the median absolute deviation (MAD) of eccentricity estimates. The procedure to compute MAD values for empirical and simulated data is described below.

For the empirical data, pRF estimates from repeated runs were grouped at the vertex level in individual subject space (fsnative). For each vertex, eccentricity values were computed from the estimated pRF center positions (*x, y*) for each run. Outliers across runs were removed using an interquartile range (IQR) criterion, where values lying more than five times the IQR beyond the first or third quartile were excluded. Such a relatively relaxed threshold was adopted to remove only extreme estimation outliers.

For each vertex *k*, the reference eccentricity value was defined as the median of the eccentricity estimates across the *in vivo* runs,

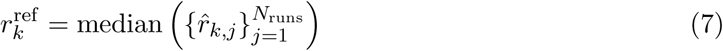

where *r̂_k,j_* denotes the eccentricity estimated for the *k*-th vertex from the *j*-th run. Residuals were then computed as

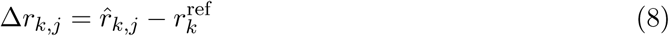

Vertices with fewer than three valid observations after outlier removal were excluded. For each remaining vertex, variability was quantified as the MAD of the residuals,

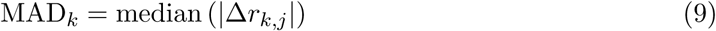

Each vertex therefore contributed a single pair *r*_*k*_^ref^, MAD*_k_*, representing its reference eccentricity and corresponding variability estimate. These pairs were subsequently assigned to fixed eccentricity bins of width 0.5*^◦^* according to *r*_*k*_^ref^. The resulting distributions of MAD values within each eccentricity bin were visualised using boxplots to characterize the variability across vertices.

For the simulated data, an analogous procedure was applied to compute MAD values. For each simulated pRF location, multiple noisy time courses (N=2000) yielded repeated pRF estimates. Eccentricity values derived from these estimates were used to compute the reference eccentricity *r*_*k*_^ref^ as the median estimated eccentricity, and residuals were computed relative to this median without reference to the known ground-truth parameters. This choice ensures consistency with the empirical data, where ground truth is inherently unknown. Importantly, it avoids biases that would arise if variability were quantified relative to ground truth, as estimation errors are not necessarily symmetric around the true parameters. The resulting pairs of reference eccentricity and MAD were assigned to the same fixed eccentricity bins as used for the empirical data.

For analyzing the correspondence of variabilities, pRF center locations and pRF sizes were sampled independently of the empirical data. Specifically, pRFs were simulated across a spatially uniform sampling of the visual field using a common range of pRF sizes at all eccentricities. Synthetic pRFs were generated using a 71 × 71 grid of pRF center locations spanning the visual field up to 24.8° (visual stimulus diameter) and 22 pRF sizes ranging from 0.1° to 2.2°, sampled at intervals of 0.1°. Each combination of pRF location and pRF size was used to generate a synthetic noise-free model time course before noise injection. Consequently, small pRFs were simulated at peripheral eccentricities and large pRFs at central eccentricities, regardless of known biological relationships between pRF size and eccentricity. For each pRF configuration, 2000 independent noisy fMRI time courses were generated at a given CNR level. This condition provides a baseline assessment of variability correspondence without incorporating empirical information regarding pRF distributions and reflects the most general use case of the simulation framework.

The resulting distributions of MAD values obtained for the simulated and empirical datasets were subsequently compared across eccentricity bins. To quantify the correspondence, the linear association between the empirical and simulated variability distributions was assessed using Pearson correlation coefficients, while the monotonic relationship was assessed using Spearman correlation coefficients.

#### 2.5.3 Analysis of RMS error across the visual field in simulated data

The procedure described in Section 2.5.2 quantifies variability using aggregated measures as a function of eccentricity. As a result, it does not provide information about the behaviour of individual pRF parameter estimates across the visual field. Therefore, pRF estimation error was additionally quantified using the root mean square (RMS) error relative to the known ground-truth pRF parameters. This provides a location-specific measure of estimation accuracy, allowing systematic evaluation of how estimation errors vary across different regions of the visual field and across pRF sizes.

For each pRF parameter *k* (i.e., horizontal position *x*, vertical position *y*, eccentricity *r*, polar angle *ϕ*, and pRF size *σ*), the RMS error was computed as

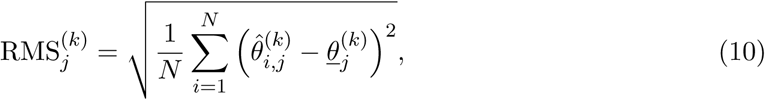

where *j* indexes the simulated ground-truth pRFs, *i* indexes the noisy realizations generated for each simulated pRF, and *N* denotes the total number of noisy realizations. Furthermore, 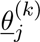 denotes the *k*-th ground-truth parameter of the *j*-th simulated pRF, while 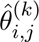 denotes the corresponding estimate obtained from the *i*-th noisy realization.

For the polar angle parameter *ϕ*, differences were computed using the circular distance to account for angular wrap-around,

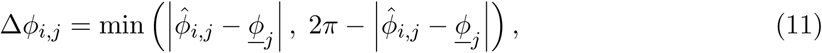

and the corresponding RMS polar angle error was computed by replacing the difference term 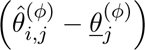 in the above equation (eq. 10) with Δ*ϕ_i,j_*. Although all calculations were performed in radians, the resulting RMS values were converted to degrees for visualization.

Only pRF estimates with variance explained (*R*^2^ *≥* 10%) were included in the analysis to reflect common thresholding procedures in pRF mapping. The goal of this analysis is to characterize how estimation error varies across spatial locations in the visual field. However, as the complete pRF parameter space comprises both spatial position and pRF size (*σ*), it cannot be visualized simultaneously. Therefore, the analysis was performed separately for each ground-truth pRF size.

For a fixed ground-truth pRF size (*σ*), RMS error values were computed for all simulated pRF center locations across the visual field. This yields a spatial distribution of estimation error for that specific *σ* value. The resulting RMS values were then arranged according to their corresponding spatial positions, forming a two-dimensional representation in which each location reflects the estimation error of a pRF centered at that position.

This procedure was repeated independently for each pRF parameter, resulting in separate spatial error maps for horizontal position (*x*), vertical position (*y*), eccentricity (*r*), polar angle (*ϕ*), and pRF size (*σ*). The maps were cropped to the circular region defined by the stimulus aperture. This analysis allows direct comparison of how estimation error varies across the visual field for different pRF parameters and ground-truth pRF sizes.

## 3 Results

### 3.1 Empirical CNR distribution

Contrast-to-noise ratio (CNR) values were computed from the empirical retinotopy dataset as described in Section 2.4.2. The empirical CNR distribution (Figure 2) was used to identify representative simulation conditions for comparison between simulated and empirical variability trends. The resulting CNR distribution was right-skewed and exhibited a median CNR = 2.36, with a substantial proportion of voxels extending toward higher CNR values. The central 50% of CNR values lay between 1.40 (Q1) and 3.63 (Q3).

**Figure 2:**
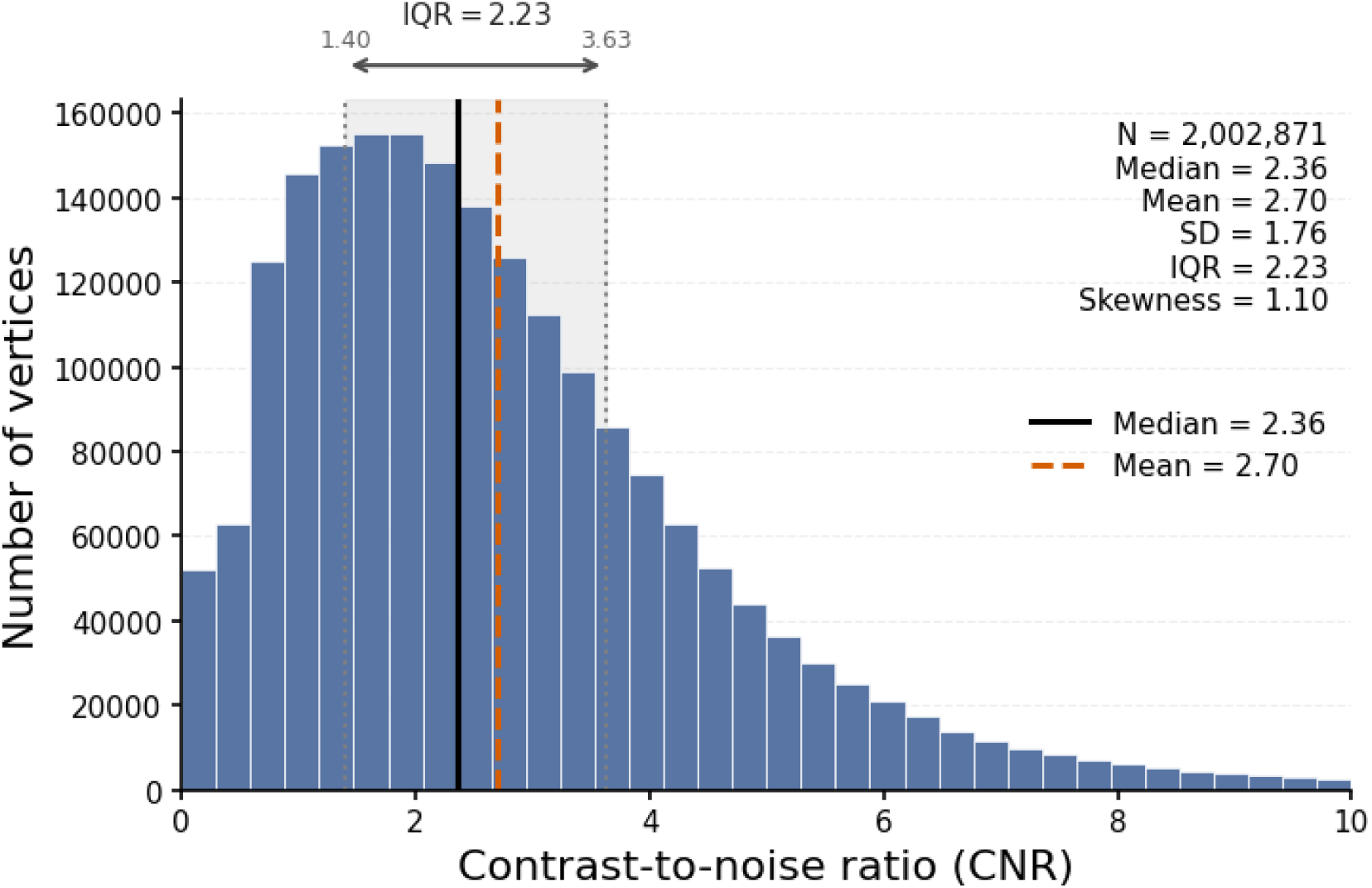
Distribution CNR values of empirical data from NYU retinotopy dataset.

Based on the empirical CNR distribution, we selected simulation conditions centered around the approximated median (CNR = 2.5) together with lower and higher neighboring conditions (CNR = 1.5 and 3.5) to examine variability trends across a representative range of signal qualities.

### 3.2 Spatial variability of simulated pRF estimates

Figure 3 presents the results of spatial variability for two representative pRF parameters as described in Section 2.5.1. For the central configuration (x=0, y=0, *σ*=1), estimated pRF centers are concentrated around the origin, with varying degrees of spatial spread, depending on the CNR level. For the peripheral configuration (x=7, y=7, *σ*=1), the estimated centers are distributed around the ground-truth location with a broader spatial spread compared to the central case. In addition, the distributions become increasingly displaced in the radial outward direction as CNR decreases.

**Figure 3:**
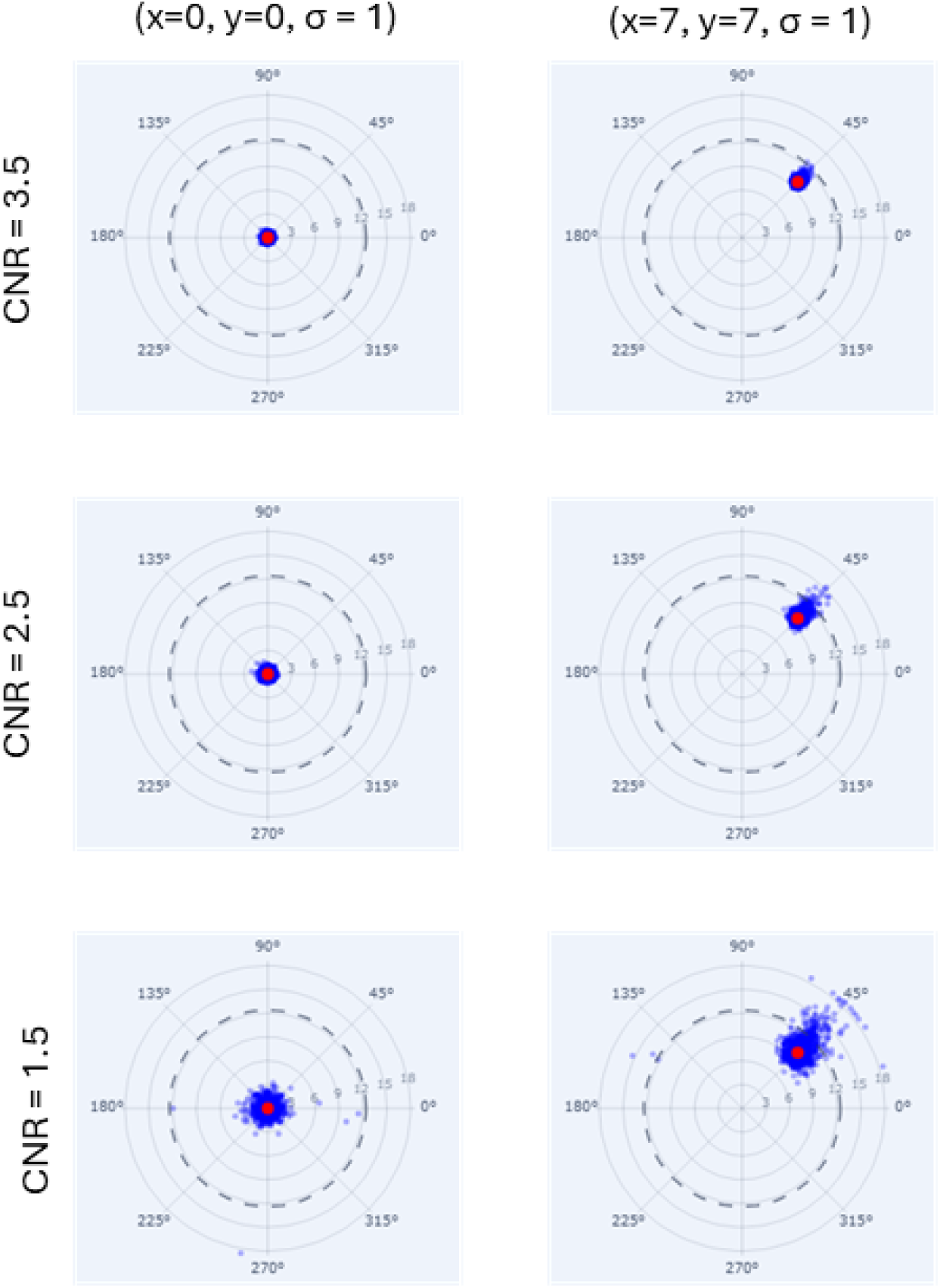
Distribution of pRF parameter estimates for simulated time courses across different ground-truth configurations and CNR levels. Each panel shows the recovered pRF centers (blue) for a fixed ground-truth location (*x, y*), indicated by the red marker. The top row corresponds to higher signal quality (CNR = 3.5), the middle row shows the results at CNR = 2.5, while the bottom row shows the results for noisier cases at CNR = 1.5. The dashed circle represents the stimulus aperture (24.8*^◦^* visual field diameter), and the outer boundary indicates the extended model search space used during pRF estimation.

Across different pRF locations and CNR levels, the spread of estimated pRF centers varies. In some cases, estimated centers extend beyond the stimulus aperture. The dashed circle denotes the stimulus extent (24.8° visual field diameter), while the outer boundary indicates the extended search space (18.5° radius, i.e., 1.5× the stimulus extent) used during model evaluation.

### 3.3 Correspondence between empirical and simulated variability

The magnitude of variability depended systematically on the simulated CNR level (Figure 4). Simulations performed at CNR = 1.5 consistently exhibited the largest MAD values across the eccentricity range, whereas simulations performed at CNR = 3.5 produced the lowest variability. The median empirical MAD values were largely bracketed by the corresponding median MAD values obtained for the CNR = 1.5 and CNR = 2.5 simulations across most eccentricity bins. The primary exception occurred in the highest eccentricity bin, where the empirical median fell below all simulated conditions. Across both empirical and simulated datasets, variability remained relatively stable over much of the visual field before increasing substantially at larger eccentricities.

**Figure 4:**
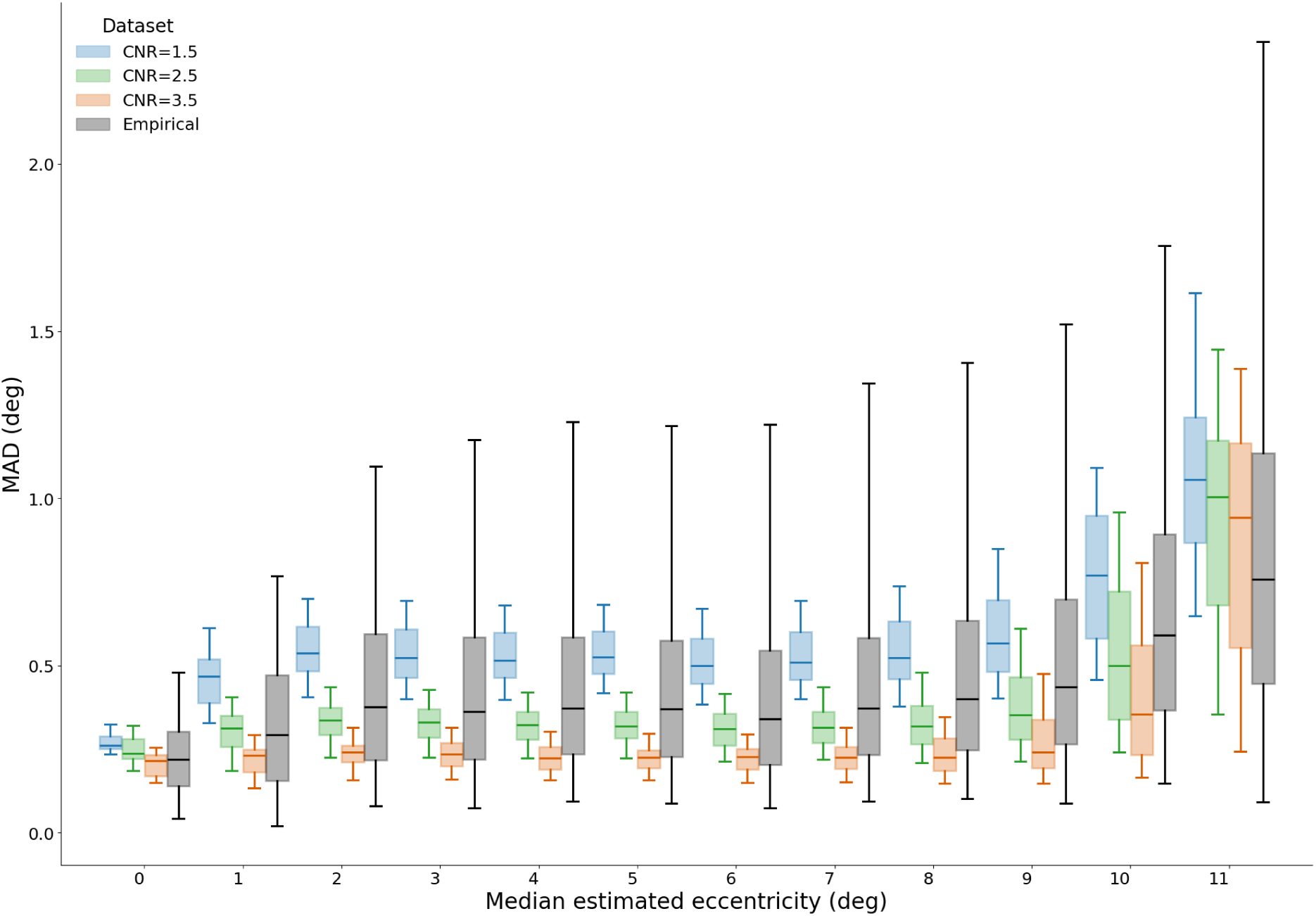
Simulated variability trends reproduce empirical observations. Median absolute deviation (MAD) of the eccentricity as a function of eccentricity for empirical pRF estimates (grey) and simulations performed using spatially uniform pRF distributions at CNR = 1.5 (blue), 2.5 (green), and 3.5 (orange). Results represent averages across bins of 0.5° width. The simulations reproduce the characteristic eccentricity-dependent increase in variability observed in the empirical data.

Quantitatively, strong agreement between simulated and empirical variability was observed for all three simulated CNR levels. Pearson correlation coefficients of median MAD values computed across eccentricity bins were 0.97, 0.95, and 0.90 for CNR values of 1.5, 2.5, and 3.5, respectively, indicating a high degree of similarity in the overall eccentricity-dependent variability distributions (Table 1). Corresponding Spearman correlation coefficients were 0.78, 0.91, and 0.89, demonstrating consistent monotonic relationships across eccentricity. The highest Spearman correlation was observed for the CNR = 2.5 condition, which also corresponds to the median CNR of the empirical dataset.

**Table 1:** Correlation coefficients between the median MAD values obtained from the empirical data and each simulated CNR condition across eccentricity bins.

| **Simulated CNR** | **Spearman $\rho$** | **Pearson $\rho$** |
| --- | --- | --- |
| 1.5 | 0.78 | 0.97 |
| 2.5 | 0.91 | 0.95 |
| 3.5 | 0.89 | 0.90 |

The boxplot representation further demonstrates that the simulations reproduce not only the overall increase in variability with eccentricity but also the magnitude of variability values. In both empirical and simulated datasets, the distributions became progressively broader towards peripheral eccentricities, reflecting greater uncertainty in pRF estimates at larger eccentricities. Although differences in the absolute magnitude of variability were observed between the simulated CNR levels, the overall eccentricity-dependent pattern remained highly consistent across all simulations.

### 3.4 Spatial distribution of RMS estimation errors

Figure 5 shows the spatial distribution of RMS errors for pRF parameter estimates obtained from simulated fMRI time courses for our chosen stimulation paradigms, i.e. NYU bar stimulus. RMS errors are displayed across the visual field for five pRF parameters: horizontal and vertical pRF center position, eccentricity, polar angle, and pRF size, as defined in Section 2.5.3. Each row corresponds to a parameter, and each column corresponds to a specific ground-truth pRF size (*σ*). The displayed columns (*σ* = 0.2, 0.4, 0.6, 1.0, 1.4, and 2.0) represent a subset of the full range of simulated pRF sizes, selected as representative examples spanning small, intermediate, and large values. All results shown correspond to a contrast-to-noise ratio (CNR) of 2.5. RMS error values vary systematically as a function of visual field location and pRF size. Spatial structure is evident in all RMS maps. For all parameters, larger ground-truth *σ* values are associated with higher RMS errors across wide regions of the visual field. Spatial patterns of RMS error are parameter-specific and stimulus-dependent.

**Figure 5:**
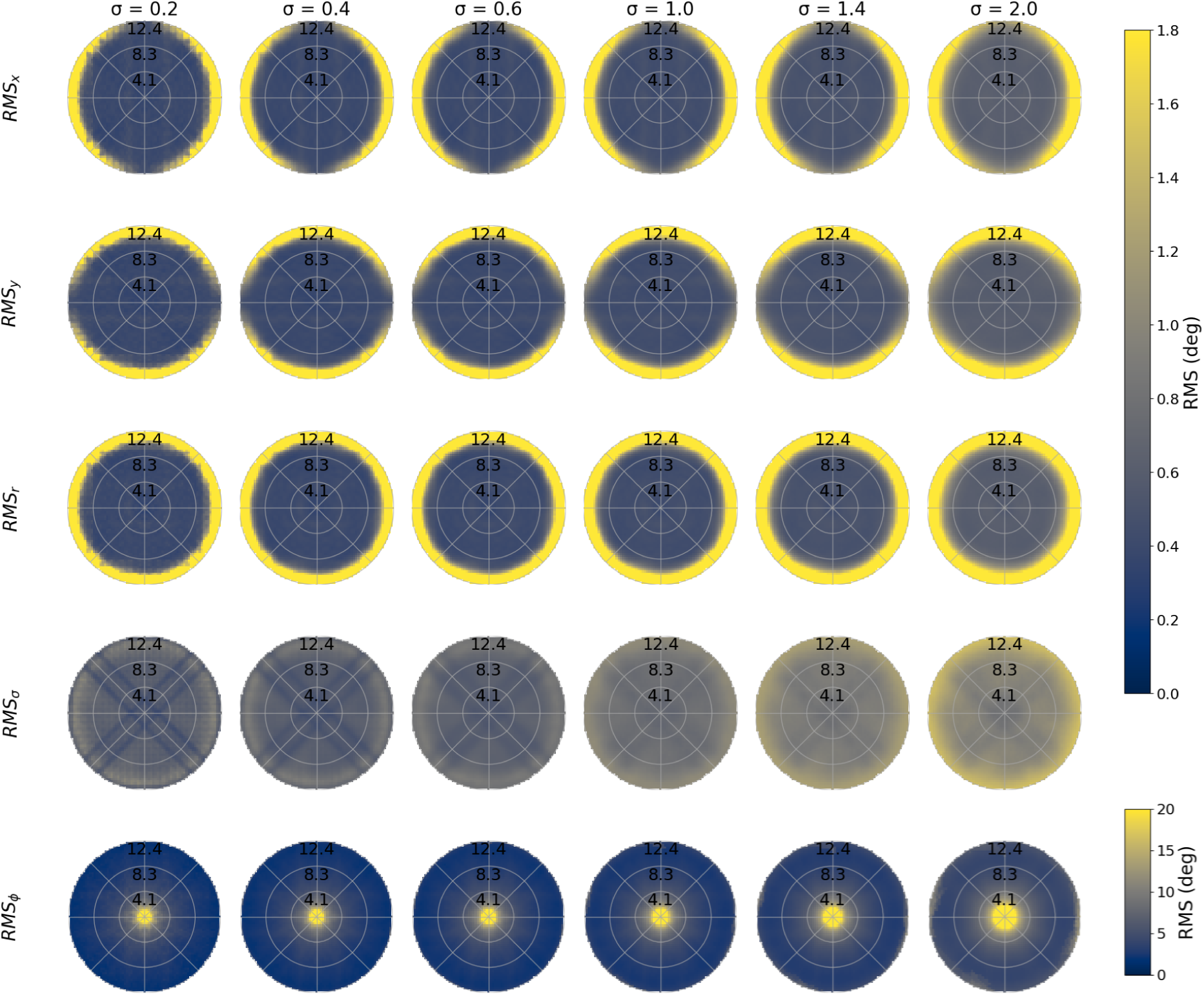
Spatial distribution of stimulus-dependent RMS pRF estimation errors. Visual field maps of RMS error for individual pRF parameters are shown for bar-type stimulation paradigms. Columns correspond to different ground-truth pRF size (*σ*) values, evaluated on the same spatial grid of simulated pRF locations for each *σ*, while rows show RMS error for pRF center position, eccentricity, polar angle, and pRF size (Section **??**). All simulations were performed under identical noise (CNR = 2.5), model, and analysis settings.

For pRF center position, RMS errors in the horizontal (x) and vertical (y) directions exhibit distinct spatial patterns that are not radially symmetric. *RMS_x_* and *RMS_y_* show anisotropic error distributions with different spatial structures along the horizontal and vertical axes. RMS errors for eccentricity increase with eccentricity. Polar angle (*ϕ*) RMS errors also show spatially structured patterns that vary with both visual field location and pRF size. However, it exhibits low error across most of the visual field and a localized increase near the central region.

RMS errors for pRF size increase with ground-truth *σ* and show only slight spatial variation across the stimulated visual field. Furthermore, the RMS error in the pRF size (*σ*) exhibits a pronounced X-shaped spatial pattern, with lower error observed along the diagonal directions of the visual field and relatively higher error in regions between these diagonals. This pattern is consistently observed across all shown ground-truth *σ* values.

Across columns, the spatial structure of RMS error remains largely consistent, while the overall magnitude varies with ground-truth *σ*. Figure 5 provides a spatially resolved overview of stimulus-dependent RMS pRF estimation errors across parameters, pRF sizes, and visual field locations under identical noise, model, and analysis settings.

## 4 Discussion

The primary objective of this study was to investigate whether large-scale simulations can reproduce the variability patterns observed in empirical population receptive field (pRF) mapping data and to identify factors that contribute to their emergence. Using the novel GEMSim-pRF framework, we performed millions of pRF simulations across a range of receptive field parameters and contrast-to-noise conditions and compared the resulting variability with repeated-measurement variability observed in the NYU retinotopy dataset. The results demonstrate that the characteristic increase in variability with eccentricity observed in empirical data was successfully reproduced in our simulations. These findings establish a close correspondence between simulated and empirical variability features and provide a foundation for simulation-driven investigation of pRF estimation behaviour and retinotopic stimulus design.

### 4.1 Large-scale simulations reproduce *in vivo* variability trends

One of the most interesting findings of this study is that the characteristic eccentricity-dependent variability observed in empirical pRF mapping results can be reproduced using large-scale simulations. Notably, this was achieved without incorporating any empirical data-based constraints. Across all investigated CNR conditions, the simulated and empirical median MAD values exhibited similar eccentricity-dependent behaviour, yielding high Pearson as well as Spearman correlation coefficient values (Table 1). One might expect that more realistic cortical pRF distributions, physiological noise sources, or other empirical constraints would be required to recover the observed variability behaviour. Instead, this analysis shows that the dominant characteristics of the empirical variability distribution emerged even in the absence of such assumptions. This suggests that the increase in variability with eccentricity is not solely a consequence of the non-uniform distribution of pRF locations across the visual field in the empirical data (e.g., due to cortical magnification [Harvey and Dumoulin, 2011]). Instead, a substantial component of the observed variability appears to arise from the interaction between stimulus geometry, signal quality, the pRF model, and the estimation procedure itself. The median of empirical MAD values is largely bracketed by simulations performed at CNR values of 1.5 and 3.5, while showing particularly strong correspondence with the median empirical CNR condition (CNR = 2.5), supporting the view that these factors alone explain the major features of the variability trend.

While the overall correspondence between simulations and empirical data was strong, a noticeable discrepancy remained at the highest eccentricity bin where empirical variability fell below all simulated conditions. At the most peripheral eccentricities, the empirical variability fell below all simulated conditions. Several factors may contribute to this behaviour. First, empirical estimates are based on comparatively few repeated measurements (up to 12 runs), and Figure 5 demonstrates that peripheral locations exhibit substantially higher estimation uncertainty. Consequently, fewer reliable estimates may remain after quality-control filtering, potentially leading to an underestimation of the empirical variability. Second, the noise model employed in the present simulations may not fully capture estimation behaviour at the visual field periphery, where additional nonlinear effects or physiological factors may become more important. Nevertheless, this local discrepancy does not alter the principal finding that both empirical and simulated data exhibit the same overall increase in variability with eccentricity.

This correspondence has a practical implication; it establishes confidence that large-scale simulations can capture variability patterns observed in real data. If the major empirical trends can be recovered under controlled settings, simulations can characterize estimation behaviour retrospectively and also predict how changes in stimulus design, signal quality, or model assumptions influence variability before empirical data are collected.

### 4.2 Role of noise in pRF estimation variability

The present findings also provide insight into the role of noise in shaping pRF estimation variability. Although the simulations employed a simple additive white Gaussian noise model, they reproduced the major variability trends observed in the empirical data. Importantly, this correspondence was not achieved merely by adding noise at any arbitrary scale to simulated time courses, but by introducing noise in a manner that preserved a consistent contrast-to-noise ratio (CNR) across different pRF configurations.

Holding CNR constant is critical because noise-free model responses vary substantially in amplitude as a function of pRF size. For pRFs modeled as 2d Gaussian, larger pRFs integrate over a broader region of the visual field and therefore generate higher-amplitude model signals than smaller pRFs. Applying identical noise amplitudes to all simulated time courses would therefore yield systematically different effective CNR values across pRF sizes and small pRFs would appear artificially more variable simply because they were evaluated at lower CNR. To avoid this confound, model time courses were normalized to remove pRF size-dependent amplitude differences before noise was added as formulated in section 2.4.1. Noise was then scaled to the target CNR, ensuring all simulated pRF configurations were evaluated under comparable signal-quality conditions. This procedure allowed the influence of pRF size, stimulus geometry, and spatial location to be examined independently from trivial differences in signal amplitude.

The empirical variability trends emerging under these controlled conditions suggest that many of the dominant features of pRF estimation variability can be simulated without using complex physiological or scanner-specific noise. A substantial variability component arises from the interaction between signal quality, spatiotemporal pattern of the stimulus, and the pRF estimation process itself. This does not imply that physiological noise, motion artifacts, or scanner-related effects are unimportant, only that they are not necessary to recover the major population-level variability trends in the empirical dataset. More sophisticated noise models may further improve quantitative correspondence, but a controlled white Gaussian noise model is sufficient to capture the dominant characteristics of pRF estimation variability.

### 4.3 Spatial structure of estimation uncertainty

In addition to our results on variability aggregated across large populations of pRF estimates, simulations also enable examining uncertainty at the level of individual pRF parameters and spatial locations. Having established a close correspondence between simulated and empirical variability distributions, we used root-mean-square (RMS) error maps computed relative to known ground-truth parameters to characterize how estimation uncertainty is distributed across the visual field.

RMS maps shown in Figure 5 reveal that estimation uncertainty is spatially structured rather than uniform, depending on both the pRF parameters being estimated and the underlying ground-truth configuration. This observation is consistent with prior work demonstrating that pRF estimation is sensitive to noise, stimulus properties, and model-stimulus interactions [Dumoulin and Wandell, 2008, Lerma-Usabiaga et al., 2020]. Furthermore, empirical studies have shown that retinotopic estimates depend on both stimulus design and visual field location [Alvarez et al., 2015, Chang et al., 2025a, Linhardt et al., 2021]. The present simulations extend these observations by providing direct access to ground-truth parameters, thereby enabling spatial patterns of estimation uncertainty to be examined explicitly.

An important observation is that the spatial structure of uncertainty differs substantially across pRF parameters. Horizontal and vertical position estimates exhibit anisotropic patterns of uncertainty, indicating direction-dependent estimation uncertainty. Eccentricity errors generally increase with distance from the fovea, while polar-angle estimates display spatially structured regions of high and low uncertainty. The localized increase in RMS for polar angle at center is expected because polar angle is undefined at the center. In contrast, pRF size (*σ*) depends globally on the underlying ground-truth size. Across all visual field locations, RMS error for *σ* increases systematically with increasing ground-truth pRF size, indicating that larger receptive fields are more prone to estimation errors.

The *σ* error maps show a pronounced X-shaped pattern that remains visible across a broad range of ground-truth pRF sizes. One plausible explanation is the potentially uneven spatiotemporal sampling properties of the stimulus. Specifically, when quantifying the number of times individual visual field locations are sampled by the bar stimulus used in the NYU retinotopy dataset [Himmelberg et al., 2021a], an X-shaped pattern emerges in which locations along the diagonal axes are sampled more frequently than surrounding regions. Denser sampling in these regions may constrain pRF size estimation, reducing estimation error. The effect may be pronounced for *σ* rather than for position parameters, potentially because pRF center locations were sampled more densely in the simulations while pRF size used a coarser parameter grid. This interpretation is consistent with the RMS structure, but disentangling stimulus sampling, parameter discretisation, and noise will require further work.

Beyond error magnitude, the simulations also revealed systematic spatial characteristics of the estimation process. For example, peripheral pRFs exhibited an increasing radial outward displacement of the estimated pRF centers as CNR decreased (Figure 3). While assessing the origin of this behaviour is beyond this study, our results illustrate that large-scale simulations can be used to identify systematic estimation characteristics that are difficult to isolate using empirical data alone.

More broadly, simulations expose behaviour that analyses of aggregated empirical datasets can not address. While empirical datasets typically provide access only to aggregate estimation outcomes, simulations show how uncertainty propagates through individual pRF parameters across the visual field. Such analyses can help identify regions of systematic estimation bias, reveal parameter-specific limitations of a stimulus design, and link population-level variability trends observed to local estimation behaviour. In this sense, the RMS maps complement the MAD-based analyses by linking global variability patterns to local estimation behaviour across the visual field.

### 4.4 Implications for simulation-driven evaluation of retinotopic stimuli

Because simulations provide access to known ground-truth parameters, they can be used to compare stimulation paradigms before empirical data are collected. For example, simulation can reveal whether a specific stimulus design systematically increases uncertainty in particular regions of the visual field, information that is difficult to obtain empirically, where the true underlying pRF parameters are unknown.

An important aspect of the present simulation framework is the normalization strategy applied. It removes the trivial amplitude scaling introduced by pRF size, so noise can be injected at consistent CNR across sizes, while preserving amplitude differences arising from the stimulus-pRF interaction. This is particularly important for future comparisons of retinotopic stimulation paradigms, as normalization schemes based on peak response amplitude would discard exactly the stimulus-dependent response differences that distinguish one paradigm from another, and that may themselves affect estimation performance.

### 4.5 Limitations and future work

Some limitations should be considered when interpreting the present findings. First, the analyses used only focused on a single retinotopic stimulation paradigm from the NYU retinotopy dataset. The large-scale simulations can reproduce the variability trends for this stimulus, but whether similar correspondence is obtained for other paradigms, such as wedge/ring stimuli [Linhardt et al., 2021] or designs incorporating cortical magnification principles [Chang et al., 2025a], remains to be tested. Second, the simulations used additive white Gaussian noise and did not explicitly model physiological fluctuations, motion artifacts, or scanner-specific noise sources [Poldrack et al., 2011]. While these are not necessary to reproduce the dominant empirical variability trends, they may contribute to finer-scale estimation behaviour and parameter-specific biases. Future work should therefore investigate whether more realistic noise models improve correspondence between simulated and empirical data. Third, empirical validation was performed using the NYU retinotopy dataset alone. Although validation using independent datasets would be desirable, assessing within-subject variability requires multiple repeated measurements of the same subject and stimulus condition, a combination rarely available. While some datasets include large numbers of subjects, they often provide only a small number of runs per subject, making reliable estimation of within-subject variability difficult. Future work would therefore benefit from independent datasets containing extensive repeated measurements acquired with different retinotopic stimulation paradigms. Despite these limitations, the present framework extends naturally to alternative paradigms and noise models, toward establishing simulation-driven evaluation as a complementary tool for optimizing retinotopic mapping.

## 5 Conclusion

Variability is a fundamental characteristic of population receptive field mapping, yet its origins have remained difficult to disentangle based on *in vivo* measurements alone. In this study, we demonstrated that the characteristic eccentricity-dependent variability observed in empirical pRF estimates can be reproduced by large-scale simulations computed under controlled noise conditions. This finding suggests that a substantial component of the observed variability emerges from the interaction between stimulus properties, signal quality, and the estimation process itself, rather than being solely determined by physiological factors. By establishing a close correspondence between simulated and empirical variability trends and by enabling direct investigation of parameter-specific uncertainty, this work provides a new framework for studying the mechanisms that shape pRF estimation. And, this framework turns retinotopic stimulus design into a testable variable: paradigms can be evaluated, and their parameter-specific biases identified, before data are collected. It can complement *in vivo* retinotopic mapping studies and support the systematic evaluation of retinotopic mapping paradigms.

## Acknowledgements

This work was supported by the Austrian Science Fund (FWF; grant https://doi.org/10.55776/P35583).

## Code Availability Statement

GEMSim-pRF will be available on GitHub as an open-source software package upon publication. It depends on GEM-pRF libraries for pRF estimation, which are also available publicly on GitHub as well as PyPI (https://pypi.org/project/gemprf/). Further instructions, usage guide and configuration tools of GEMSim-pRF and GEM-pRF are part of the documentation website (https://gemprf.github.io/).

## AI Statement

The authors used AI-based tools (ChatGPT and Grammarly) to assist with language editing and software development. All generated content was carefully reviewed and verified by the authors, who take full responsibility for the final manuscript.

